# Genetic Diversity and Coexistence of *Babesia* in Ticks (Acari: Ixodidae) from Northeastern China

**DOI:** 10.1101/469833

**Authors:** Jia-Fu Jiang, Bao-Gui Jiang, Ting-Ting Yuan, Michael Von Fricken, Na Jia, Rui-Ruo Jiang, Yuan Zhang, Xin-Lou Li, Yuan-Chun Zheng, Qiu-Bo Huo, Yi Sun, Wu-Chun Cao

**Affiliations:** State Key Laboratory of Pathogen and Biosecurity, Beijing Institute of Microbiology and Epidemiology, Beijing, P. R. China; George Mason University, Dept. of Global and Community Health, Fairfax, VA, USA; Mudanjiang Forestry Central Hospital, Mudanjiang, P. R. China

**Keywords:** *Babesia*, ticks, diversity, Coexistence, China

## Abstract

**Background:** Babesiosis is an emerging zoonosis in humans with significant and increasing health burden in China. A few systematic reports on *Babesia* spp. was involved with ticks, especially in the human babesiosis endemic areas.

**Methods:** The ticks were collected from 30 individual waypoints along 2.0 km transects in two recreational forested areas in Northeastern China. Then we screened them for *Babesia* spp. infection by amplifying the partial 18s rRNA gene with subsequent sequencing. Multivariate logistic regression analysis was used to access the association between infections and some related risk factors. The cluster analyses were performed using SaTScan v6.0 Software for identifying the geographic cluster of the positive samples in ticks from each waypoint.

**Results:** A total of *Ixodes persulcatus* (n=2380) and *Haemaphysalis concinna* (n=461) ticks were collected. The 0.97% of *I. persulcatus* ticks were infected with five *Babesia* species, including *B. bigemina* (n=6), *B. divergens* (n=2), *B. microti* (n=3), *B. venatorum* (n=11) and one novel strain HLJ-8. Thirteen (2.92%) *H.concinna* ticks contained *B. bigemina* (n=1), *B. divergens* (n=1), three genetic variants of *Babesia* represented by HLJ-874 which was closely related to *Babesia* sp.MA#361-1, and eight other *Babesia* variants represented by HLJ242 which were similar to *B.crassa*. Each study site had 5~6 different *Babesia* spp. One waypoint was more likely to yield *B.venatorum* (relative risk=15.36, *P*=0.045) than all other waypoints.

**Conclusions:** There exists a high genetic diversity of *Babesia* spp. across a relatively small sampled region. Further study is needed to understand the risks these variants pose for human health.

**Author Summary:** Babesiosis is the subject of increasing interest as an emerging zoonosis in humans with significant and increasing health burden of the disease at recently. In China, many probably human babesiosis cases who had a history of recent tick bite were found in Lyme endemic area in Northeastern China, where the prevalence of Babesia parasite in the ticks still was far underestimated. In the present study, we conducted a field survey for ticks to identify diversities and complexity of babesia, and then to assess the risk of human babesiosis, by means of a three years longitudinal study that mapped the location of the ticks tested positive for Babesia spp. at two forestry areas with a heavy burden of tick-borne pathogens. We firstly presented the prevalence of Babesia spp. especially the genetic diversities and coexistence of seven Babesia spp. including 2 novel species or variants at one small scale “natural foci” in northeastern China. This work is useful to understand the complexity of Babesia pathogen in China, and how the Babesia perpetuates over the long term in the environment, as well as potential risks for human health.

## Introduction

Babesiosis, caused by infection with intraerythrocytic parasites of the genus *Babesia*, is one of the most common infections of free-living animals globally[1]. Cases of babesiosis have been reported in Europe, Africa, Australia, South America and Asia [1], with 1,092 confirmed and probable cases reported across seven states of the United States in 2011 [2]. More than 100 *Babesia* species have been documented in a wide variety of wild and domestic animals, some of which are known to infect humans [3]. In America, *Babesia microti* is the most important causative agent of human babesiosis, followed by sporadic cases of *B. duncani and B. divergens*–like infections[1]. In Europe, the primary agent is *B. divergens*, but a small number of *B. microti* and *B. venatorum* (formerly known as Babesia sp. EU1) patients have also been reported in Austria, Germany, and Italy [4]. In Asia, *B. microti*–like organisms have been reported to cause human infection in Japan, Korea, and China[5-8]. In China, many *Babesia* parasite isolates, including *B. bovis, B. equi, B. caballi*, and *B. gibsoni have been* documented in domestic animals[9, 10]. A few studies were about human cases of babesiosis in China[8, 11-15] warranting additional efforts to study this neglected disease.

Besides via blood transfusion, transmission of *Babesia* occurs primarily through ticks. *B. microti is transmitted by I. ovatus and I. persulcatus in Asia*[6, 7]. Additionally, *I. persulcatus, I. ricinus* and *H. japonica* in Far East Russia were also found to be mainly infected with *Babesia microti* [7, 16, 17], *B. venatorum* [16], *B. divergens*, and *B. capreoli* [17]. In China, reports of *Babesia* spp. were attributed to ticks, and confirmed through isolation and genetic sequencing[18, 19]. In recent years, cases of probable or subclinical human babesiosis in patients reporting a recent tick bite have been recorded in Lyme disease-endemic areas of Northeastern China [14, 15]. The similarity in symptoms and potential for misdiagnosis present the possibility that the prevalence of *Babesia is underestimated. In these areas*, there is still a lack of detail information on pathogenic or non-pathogenic *Babesia* spp. in ticks and humans.

This three-year longitudinal study maps the location of ticks that tested positive for *Babesia* spp. in two forested areas known to be endemic with other tick-borne pathogens, like Lyme disease and tick-borne encephalitis virus. We conducted a field study to identify the prevalence and molecular diversity of *Babesia* in sampled ticks.

## Material and methods

### Ticks collection

Host-seeking ticks were collected between April to July from 2010 to 2012 by flagging vegetation in 30 individual waypoints along about 2.0 km transect respectively at two recreational forest areas. Dashigou scenic area(E129°11′10″, N44°58′32″)and Heiniubei scenic area (E129°27′31″, N45°2′07″), are both located in the Daxinganling Mountains near Mudanjiang city in Heilongjiang province, about fifty kilometers apart. The spatial distribution of sample sites and collected ticks were mapped in ArcGIS 9.3 software (ESRI Inc, Redlands, CA, USA). Identification of tick species was done according to the key morphologic features described by Teng and Jiang[20].

### DNA extraction and Amplification of *Babesia* parasites

Each tick was crushed individually in a sealed micro-centrifuge tube with Buller Blender Homogenizer (Next Advance Inc., NY, USA). DNA was then extracted from tick homogenate using the Tissue DNA Extract Kit (Tiangen Biotechnique Inc., Beijing, China) according to the manufacturer’s instructions. To ensure amplifiable DNA was extracted and conformed to the identified tick species, PCR for mitochondrial 16S rRNA employing tick-specific primers (5’ CCGGTCTGAACTCAGATCAAG-3’, and 5’ CAATGATTWTTTAAATTGSTGTGG-3’)[21] was performed. PCR targeting the specific fragment encoding the partial 18S rRNA gene was performed with modified primers PIRO-A and PIRO-B to screen for *Babesia* spp. infection[22]. For *Babesia* spp. with unclear taxonomy based on the gene segment above two pairs of designed primers (5’ GAAACTGCGAATGGCTCATTACAACA-3’ and 5’ CAACCGTTCCTATTAACCATTA-3’; 5’ TAATGGTTAATAGGAACGGTTG-3’ and 5’ CTACGGAAACCTTGTTACGACTT-3’) were used for the amplification of near-entire sequences of the 18S rRNA gene for most of *Babesia* spp. in Asia. The other two pairs of designed primer (5’ CTCGCGAATCGCAATTTA-3’ and 5’ ACAGACCTGTTATTGCCTTAC; 5’-AAATTAGCGAATCGCATGG-3’ and 5’-ACAG ACCTGTTATTGCCTTAC-3’) were used for amplification of entire sequences of the 18S rRNA of *B. microti*. The 30 µl reaction mixture used 2 µl of genomic DNA as template, 1 U Taq polymerase (Takara), 3 µl of 10× PCR reaction buffer, 3 µl of 10 mM MgCl_2_ (final concentration 1.5 mM), 0.6 µl of 2.5 mM dNTPs mixture (final concentration 0.5 mM), and 1 µl of each primer (final concentration 0.4 mM). A three-step thermal cycling program was used to amplify the target gene fragment. The amplified product was visualized in UV light. To avoid possible contamination, DNA extraction, the reagent setup, amplification, and agarose gel electrophoresis were performed in separate rooms, and negative control samples (distilled water) were included in each amplification.

### Sequencing and phylogenetic analysis of the *Babesia* parasites

The amplification product was purified by TIANgel Mini Purification Kit (Tiangen Biotechnique Inc.) following the manufacturer’s instructions. The short strands below 500 bp were sequenced directly. The long purified DNA fragments were cloned into the plasmid pGEM-T vector and transformed into competent cells (XL1-blue *Escherichia coli*). The recombinant plasmids were then extracted and purified. The nucleotide sequences of the plasmid inserts were determined using an automated DNA sequencer (3730 DNA Sequencer, Applied Biosystems). The alignment and assembly of sequences, calculation of phylogeny distances, and construction of phylogenetic trees were performed using MEGA 5.0 software [23].

### Risk factors analyses

Univariate logistic analysis was conducted to determine the associations between the prevalence of *Babesia* spp. and sample sites, vector species, vector gender, forest stand composition, niche, elevation, location of hill, shape, direction and degree of slope, dominant/secondary tree, dominant/secondary shrub and herbaceous plant grass. A *P* value < 0.05 was considered statistically significant. Odds ratios (ORs) were estimated by comparing infection status with suspected risk factors. Multivariate logistic analysis was then performed using variables with *P* values < 0.10 from the univariate analyses as covariates.

### Spatial cluster analyses

In order to identify the geographic cluster of the positive samples in ticks from each sample sites, cluster analyses were performed using SaTScan v6.0 Software 2005 (http://www.satsan.org). SaTScan uses a circular window that moves through space to identify clusters. The window varied in size for up to 50% of the population tested, allowing for the identification of small and large clusters. A likelihood ratio test was conducted to determine whether there was an elevated rate of ‘cases’, or infected ticks, in one area compared to surrounding areas. Significance (set a priori at *P*<0.05) was then calculated using Monte Carlo replicates.

## Results

### The prevalence of *Babesia* spp. in ticks

A total of 2380 questing adult *I. persulcatus* and 461 *H. concinna* ticks were collected over three years (2010–2012) from a total of 60 waypoints in our two field study sites. A minimum of 791 ticks was examined each year (Table 1). The annual prevalence of ticks testing positive by PCR for *Babesia* DNA ranged from 1.02% to 1.62% (Table 1). There was no significant difference between the prevalence in different years. The prevalence of *Babesia* between the two regions was significant (*P*=0.04), with 1.74% (24/1383) positive in Heiniubei and 0.82% (12/1458) in Dashigou. The overall prevalence in *I. persulcatus* ticks was 0.97 % with 23 positive ticks, 10 male and 13 female. Thirteen (2.81%) *H. concinna* ticks, including eight male and five female, contained *Babesia* DNA. The *Babesia* infection rate between the two tick species was significant (*P*=0.001), but no association was observed by gender (Table 2).

**Table 1.** Population Index of collected ticks and Prevalence of the agent of Babesia spp. in ticks at two study sites from 2010-2012.

| Year | Heiniubei (S1) |  |  | Dashigou (S2) |  |  | Total |  |  |
| --- | --- | --- | --- | --- | --- | --- | --- | --- | --- |
|  | Number | Prevalence | Population Index* | Number | Prevalence | Population Index | Number | Prevalence | Population Index |
| 2010 | 234 | 1.71% | 13.0 | 689 | 0.88% | 28.7 | 923 | 1.10% | 22.0 |
| 2011 | 591 | 2.03% | 16.4 | 536 | 1.16% | 29.7 | 1127 | 1.62% | 20.9 |
| 2012 | 558 | 1.43% | 18.6 | 233 | 0 | 7.7 | 791 | 1.02% | 13.2 |

\*Average number of ticks collected per hours each person.

**Table 2.**
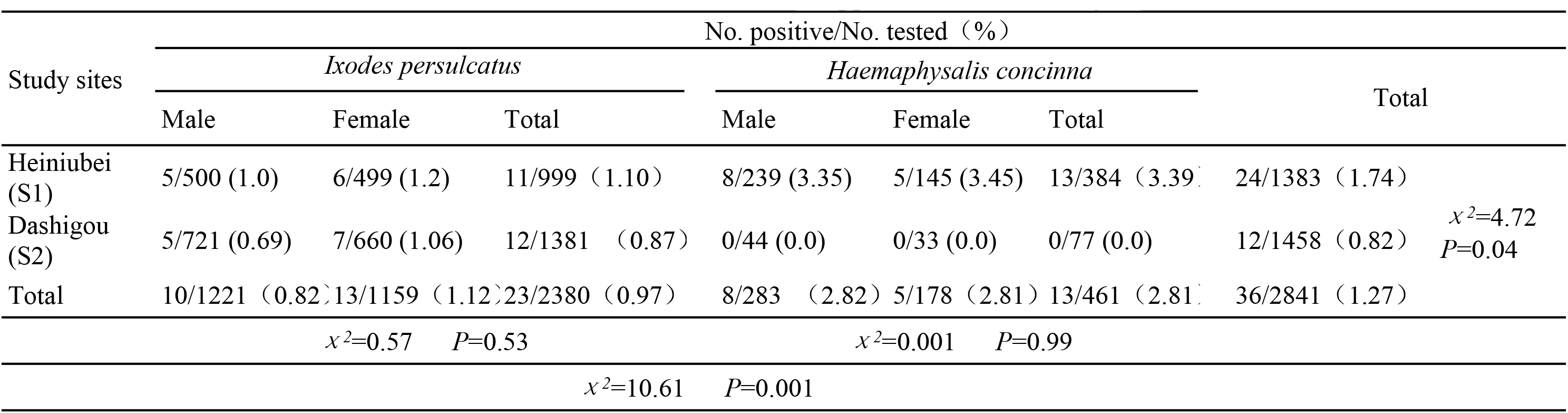
Results of PCR for *Babesia* spp. of ticks at two study sites.

### Sequence comparison and diversities of *Babesia*

Based on the ~400 bp recovered from 36 positive samples, through the alignment and BLAST for highly similar nucleotide sequences GenBank (PubMed website: www.ncbi.nlm.gov/BLAST/) showed that the sequences from two *I. persulcatus* and one *H. concinna* shared 100% identity and was most closely related to *B. divergens* isolate Nov-Ip316 (GU057385) from *I. persulcatus* in Novosibirsk, Russia. The sequences from six *I. persulcatus* and one *H.concinna* shared 100% identity and 98% with *B. bigemina* strain 563 (HQ840960) respectively. The sequences from 11 *I. persulcatus* ticks represented by HLJ223 (accession No. JQ993246) shared 100% identity (406 bp) with *B. venatorum* from *I. persulcatus* in Novosibirsk, Russia (GU734773) and patients from Austria and Italy (AY046575). The sequences recovered from three *I. persulcatus*, HLJ407, HLJ247 and HLJ72 (JQ993429) shared 100% identity (435bp) with *B. microti* from *I. persulcatus* at Novosibirsk in Russia (GU057383). One *I. persulcatus* (HLJ8, accession No. KF582564) had high variation (more than 8% difference) with our study samples and share 97% identities with *Babesia motasi* (AY260180). Three *H. concinna* represented by HLJ874 (accession No. JQ993427) was most closely related to *Babesia* sp. MA631 isolated from raccoons in Japan. The sequences from eight *H. concinna* ticks represented by HLJ242 had 2~7 bp differences at position 200~250 nt from *Babesia crass* detected in sheep in Iran (AY260176).

Our study found both *I. persulcatus* and *H. concinna* ticks can carry four and five species of *Babesia* spp. respectively. The former included *B. bigemina* (n=6), *B. divergens* (n=2), *B.microti* (n=3), *B. venatorum*-like protozoan (n=11) and one variation HLJ-8; the latter included *B. bigemina* (n=1), *B. divergens* (n=1), *Babesia crass*-like (HLJ-242) protozoan (n=8) and one kind of unidentified *Babesia* sp. HLJ-874-like protozoan (n=3) and The species of *Babesia* carried by these two ticks had significant differences (*P*=0.001), and showed high diversity, and to some extent vector tropism. Also, the two sites studied showed significant differences (*P*=0.01) in the distribution of different kinds of *Babesia* spp. (Table 3). Through phylogenic analysis, the sequences from our study formed a closely related clade with *B. venatorum* (Babesia sp. EU1), *B. microti*, *Babesia* crass-like, *Babesia* sp. MA361, *B. bigemina* and *B.divergens*, respectively (Figure 1A).

**Table 3.** Results of infected ticks by different *Babesia* species.

| Sample sites <sup>a</sup> Tick species <sup>b</sup> |  | No. tested | No. positive |  |  |  |  |  |  |  |
| --- | --- | --- | --- | --- | --- | --- | --- | --- | --- | --- |
|  |  |  | *Babesia bigemina* | *Babesia divergens* | *Babesia venatorum* | *Babesia microti* | *Babesia sp* HLJ-8 | *Babesia sp.* HLJ-874 | *Babesia sp* HLJ-242 | Total |
| Heiniubei | *Ixodes persulcatus* | 999 | 3 | 0 | 8 | 3 | 0 | 0 | 0 | 11 |
|  | *Haemaphysalis concinna* | 384 | 1 | 1 | 0 | 0 | 0 | 3 | 8 | 13 |
| Dashigou | *Ixodes persulcatus* | 1381 | 3 | 2 | 3 | 3 | 1 | 0 | 0 | 12 |
|  | *Haemaphysalis concinna* | 77 | 0 | 0 | 0 | 0 | 0 | 0 | 0 | 0 |
| Total |  | 2841 | 7 | 3 | 11 | 3 | 1 | 3 | 8 | 36 |

<sup>a</sup> $\chi^2=18.48$ $P=0.01$ ; <sup>b</sup> $\chi^2=60.50$ $P=0.001$ ;

**Figure 1.**
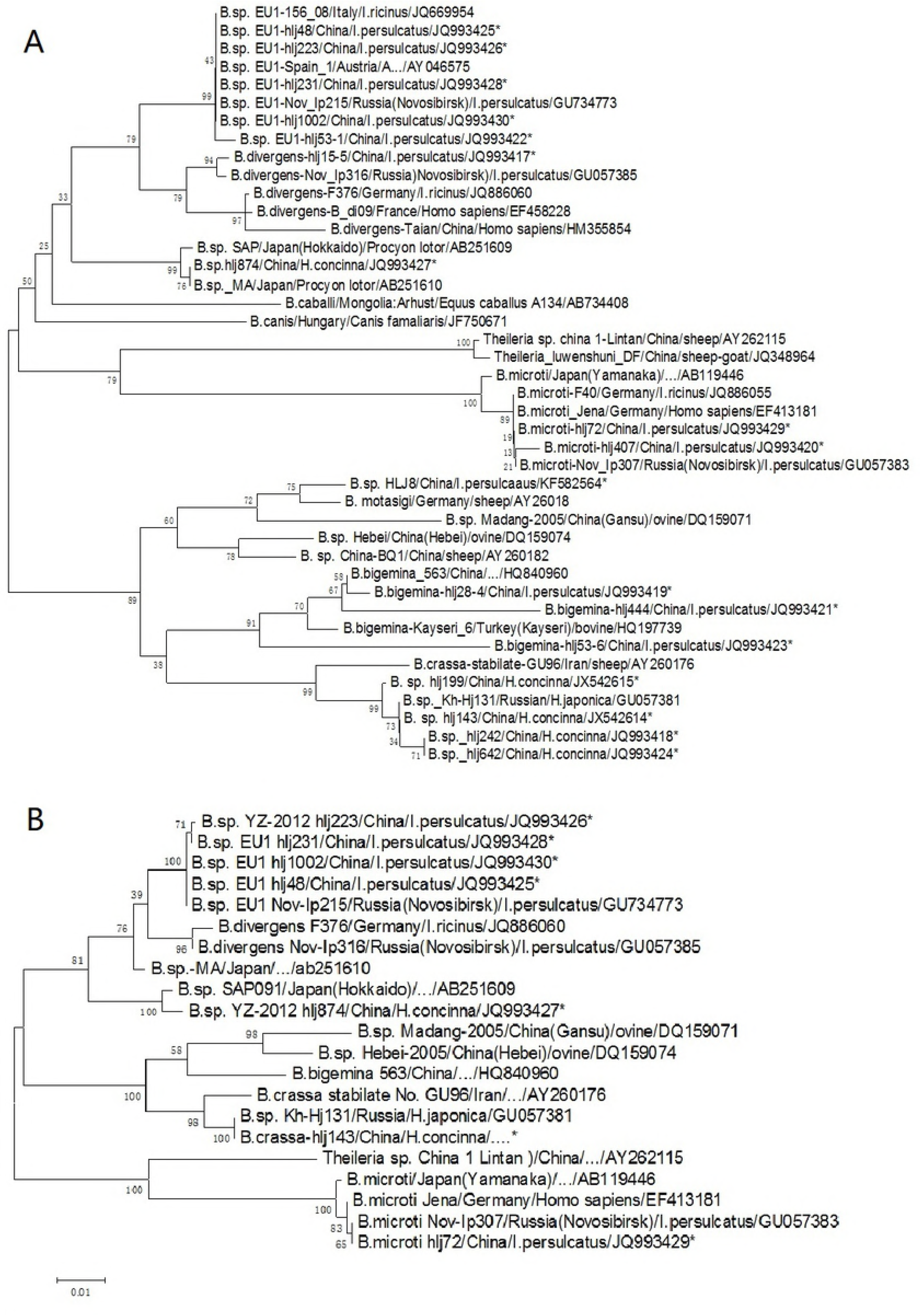
Phylogenetic tree based on partial (406~435 bp) (A)18s rRNA sequences of *Babesia* spp., and near entire 18s rRNA (B), obtained by using neighbor-joining method with Kimura 2-parameter analysis and bootstrap analysis of 1,0000 replicates. Numbers on the branches indicate percentage of replicates that reproduced the topology for each clade. Parentheses enclose GenBank numbers of the sequences used in the phylogenetic analysis. The asterisk (*) indicates novel sequences obtained from tick in this study. Scale bar indicates number of nucleotides per 1,000 bp *Babesia spp*.

For the *Babesia* spp. samples which had unclear taxonomy and three *B. microti*-like samples which had divergence from other known species in this study, nearly the entire 18s rRNA sequence of *Babesia* was analyzed. The 1600~1688 bp sequences of *Babesia* recovered from the HLJ48, HLJ223, HLJ231 and HLJ1002 were identical to each other, and had 2 nt difference at the 1277 bp and 1443 bp position with *B. venatorum* from patients in Austria and Italy (AY046575). The 1605 near-entire *rrs* sequences of *Babesia* from the HLJ143, HLJ199 and HLJ242 had highest homology (97.5%) with *B. crass* (AY260176) in the GenBank. However, through analysis with Mega 5.0 software, there was only 2 nt difference among 1100 bp with those of *Babesia* sp. Kh-hj131, which had no whole genome sequences available in GenBank. The sequences of HLJ72 and HLJ874 also showed the highest homology with *B. microti-*Nov Ip307 *and Babesia* sp. MA361 respectively. HLJ72 (JQ993429) also shared 100% identity with *B. microti* from *I. persulcatus* at Novosibirsk in Russia (GU057383). The phylogenetic tree also showed the same topological graph (Figure 1B) as the results with those based on 400-bp sequences.

### Risk factors

Univariate logistic regression analysis for sample sites, vectors species, and shape of slope elevation, showed significant correlation with *Babesia* prevalence in ticks. Ticks in Heiniubei sites (OR = 2,13, 95% CI =1.06–4.27), and *Haemaphysalis concinna* (OR = 2.97, 95% CI =1.50–5.91), had higher infection rates than those of other corresponding factors, respectively. Through stratified analysis by site factor, there were no significant factors associated with the *Babesia* infection in the ticks at Dashigou sites. However, for Heiniubei sites, the forest stand composition, location in the hill, shape of slope, dominant grass and endemic tick species were significantly correlated with *Babesia* infection in ticks. *H. concinna* (OR =3.15, 95% CI =1.40–7.09), ticks in the concave slope (OR =8.59, 95% CI =1.02-72.31), in either wild weeds *Equisetum ramosissimum* (OR =10.27, 95% CI =1.85–56.91) and *Adonis amurensis* (OR = 6.15, 95% CI =1.23–30.73), had higher infection rate than those of other corresponding factors (Table 4). Multivariate logistic regression analysis indicated that tick species (*H. concinna*), shape of slope (concave slope), and land-dominant grass (*Equisetum ramosissimum*) were significantly associated with the prevalence of *Babesia* infection in ticks (Table 5).

**Table 4.**
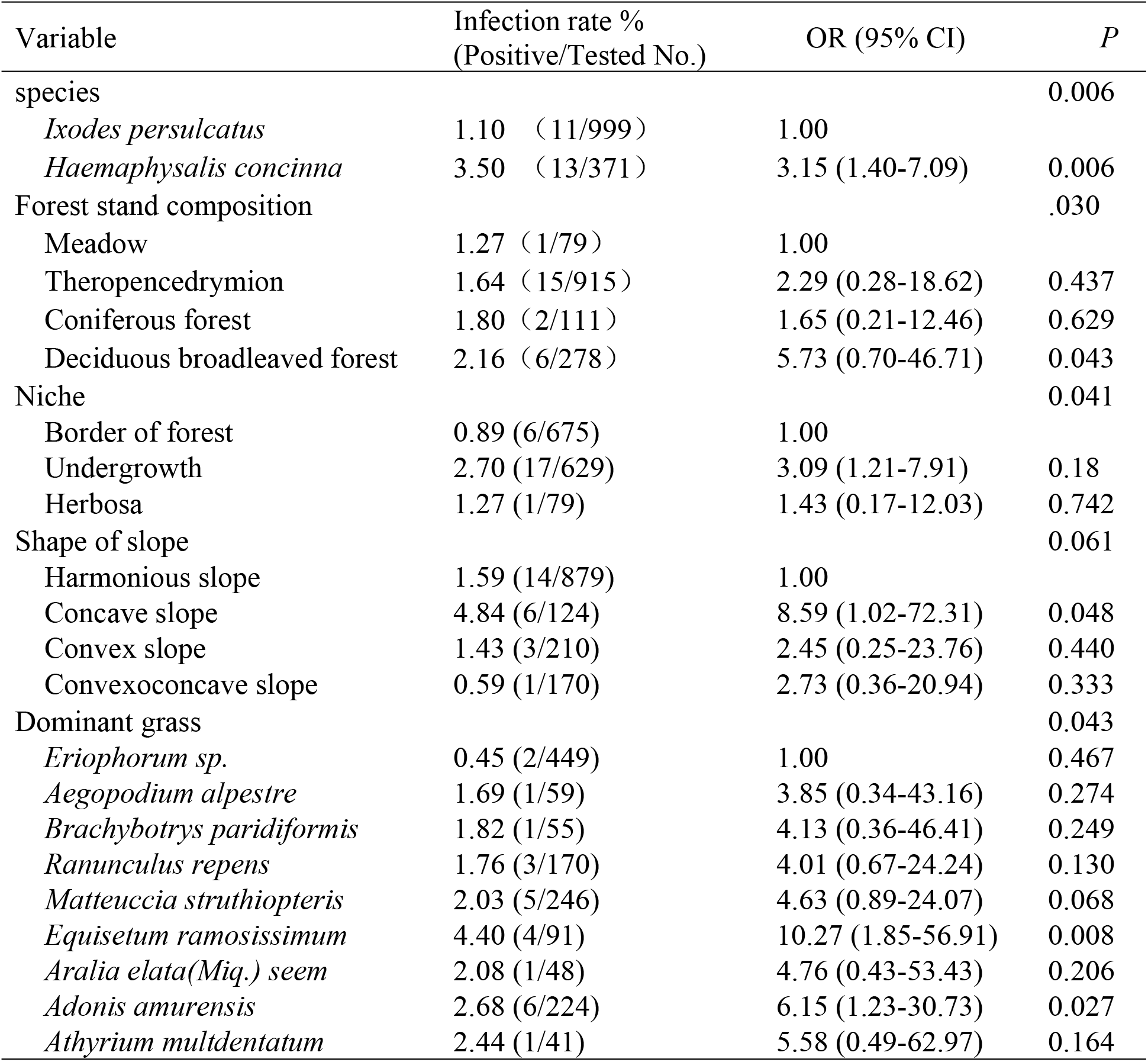
Results of univariate logistic regression analysis for Babesia infection in ticks from site Heiniubei at Northeastern China.

**Table 5.** Results of Multi-variate logistic regression analysis for Babesia infection in ticks from site Heiniubei at Northeastern China.

| Variable | OR (95% CI) | *P* |
| --- | --- | --- |
| Species | 3.54 (1.57-8.24) | 0.003 |
| Concave slope | 46.01 (1.38-152.9) | 0.03 |
| *Equisetum ramosissimum* | 6.95 (0.75-63.75) | 0.046 |

### Spatial distribution of collected ticks and positive samples

The spatial distribution of total collected ticks was mapped (Figure 3) in each waypoint sample site, which varied from 24 to 89 during 3 years. We overlapped the positive samples by using different colors and shapes which represent the collected time. One waypoint in site S2 (Dashigou, Figure 2A) and three waypoints in site S1 (Heiniubei Figure 2B) had positive samples for two continuous years. The remaining fifteen positive waypoints were all randomly distributed over different years, with infection rate varying between 0 to 4.26% (mean infection rate = 0.79%) at site S2, and 0 to 8.33% (mean infection rate = 1.68%) at site S1. However, the cluster analysis for data of site S1 by different *Babesia* species or variants demonstrated that there were significant clusters in each field, indicating a relative cluster distribution of positive ticks for *B. venatorum* and HLJ242-like variants (Figure 3). Positive ticks collected in the cluster (waypoint) were 15.36 times (relative risk = 15.36, P=0.045) more likely to be infected with *B. venatorum* than those collected elsewhere in the transect sample site.

**Figure 2.**
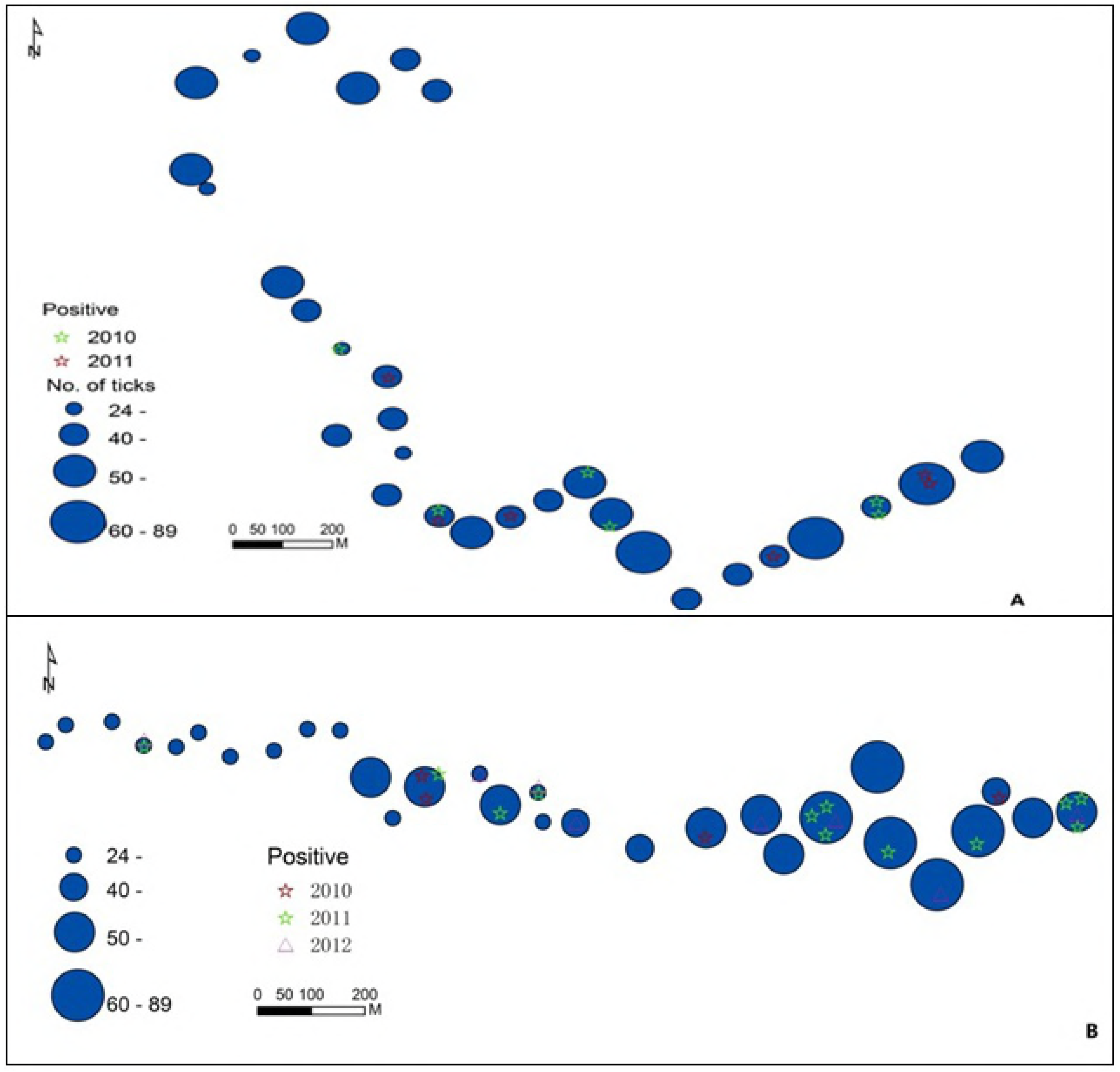
GIS mapping of the field site, total number of ticks collected at each 30 individual waypoints and distribution of positive ticks in Dashigou (A) and Heiniubei (B) respectively, 2010-2012. (The size of the dot at each waypoint is proportional to the number of ticks at each site. Each ticks that tested positive for *Babesia* spp. DNA by PCR from each waypoint was mapped).

## Discussion

Here we examined *Babesia* spp. infection of 2841 ticks (2380 *I. persulcatus* and 461 *H. concinna*) through amplification and sequencing of partial 18S rRNA. This is a more reliable method to distinguish the deeper branches among interspecies than through the partial cytochrome b gene and internal transcribed spacer (ITS) gene [24]. Although the prevalence (1.27%) of *Babesia* in not high, at least seven *Babesia* species coexisted in these relatively small areas. This study indicates that the discovered agents show genetic diversity, complexity, and are more widely distributed among ticks in Northeastern China than previously recognized. As the interface between humans and wildlife increases, humans may come into contact with these tick species more often, introducing greater risks to human health.

In our study, 0.46% (11/2380) of the examined *I. persulcatus* ticks were infected by *B. venatorum* (formerly called *Babesia* sp. EU1). This *Babesia* species was first characterized from patients in Europe [4], and has been identified in roe deer, wild *cervids, migratory birds*, and host-seeking *I. ricinus* and *I. persulcatus*[16, 17, 25-27]. To our knowledge, it has been poorly investigated in ticks, rodents, livestock and humans in China. We confirmed the presence of *B. venatorum* and the four obtained entire sequences showed 99.9 % similarity to *B. venatorum* isolate 7627 from deer and strain BAB20 from patients in Europe. Our results also showed that the *B. venatorum* is a predominant *Babesia* species in *I. persulcatus* in these sampled areas. Our finding is identical to the main species of *Babesia* recovered from the patients in these regions[14].

*B. divergens* was detected in two *I. persulcatus* and one *H. concinna*. The recovered sequences showed 100% homology to those reported from an infected *I. persulcatus* at Novosibirsk in Russia, which is geographically far from northeastern China[28]. This is also the first detection of *B. divergens* in ticks collected in China. Recently *B. divergens* protozoan was found in two anemic patients in Shandong province, which is far from northeastern China, indicating that this isolate may be pathogenic to humans [13]. The vector of transmission in this case is also unclear, as there are no reported *I. persulcatus* and *H. concinna* ticks in Shandong.

*Babesia microti* was identified in 0.13% (3/2380) of examined *I. persulcatus* in our study sites. This infection rate is low compared with those in Suifenhe city, which is roughly 200-kilometers from our study sites[18]. Various studies in Europe also show *B. microti* present in *I. ricinus* with prevalences reported between 1.3% to 11.1% in Poland[29, 30], 10% in Slovenia (31), and 4% in Switzerland (32). In Asia, *B. microti* was detected in 12.3% of *I. ovatus* and1.4% of *I. persulcatus* in Japan[2], and 0.29% (1/347) from Novosibirsk and 1.3% (1/77) in Khabarovsk Territory, which is near our study sites[28]. This suggests a wide distribution of *B. microti* in ticks. Our study also showed the predominant presence of *B. bigemina*, which was detected in 0.25% (6/2380) of *I. persulcatus* and 0.21 % (1/461) of *H. concinna*. Sequenced isolates shared 98% identities with *B. bigemina* strain 563 from *Bubalus babalis*. This organism is thought to be transmitted by *Boophilus microplus, Rhipicephalus haemaphysaloides*, and *H. longicornis* ticks, and responsible for disease in dairy cattle, buffaloes and dogs in southern China[31]. However, little information on *B. bigemina* in northeastern China is known. As *I. persulcatus* and *H. concinna* often bite humans, the pathogenicity of this organism for humans is an important issue warranting further investigation.

Three genetic variants of *Babesia* strico lato, represented by HLJ-874 strain, were closely related to *Babesia* sp. Ma 361 isolated from a feral raccoon in Japan[26]. The other genetic variants recovered from eight *H. concinna* ticks representing HLJ-143 strain were most similar to *Babesia* sp. Hj131 found in *H. japonica* tick from Khabarovsk Territory in Far East Russia based on partial 18S rRNA gene. This was similar to the human *B*. crassa-like pathogen (99%)[15] and ovine pathogen *B*. crassa (97%) on the basis of entire 18S rRNA gene sequence[28]. One genetic variant of *Babesia* strico lato, HLJ-8 strain recovered from *I. persulcatus* was closely related to *B. motasi* (AY260180) isolated from a feral raccoon in Japan[26]. It suggests that our HLJ-874- and HLJ-143-like *Babesia* spp. are new species or recordings in China. These results may indicate at least two yet un-described *Babesia* species exist within these small natural foci.

*I. persulcatus* was infected with four *Babesia* species (*B. bigemina, B. divergens, B. microti* and B. venatorum) and H. concinna was *infected with three, possibly four, Babesia species (B. bigemina*, *B. divergens* and 2 unidentified *Babesia* spp.). These disparate species may reflect differences in vector competence for each kind of *Babesia* parasite. It is unclear if these *Babesia* spp. can be efficiently transmitted by ticks, moreover, the prevalence of *Babesia* spp. infection in nymphal ticks requires further collection and analysis.

This study also showed the heterogeneity in the spatial distribution of *Babesia* spp. in ticks at small scale. It is not surprising that forest stand composition features, niche, shape of slope, and dominant grass may have influenced tick density and distribution, as specific environmental habitats may be more suitable for certain species of ticks. The two predominant ticks in these areas have relatively different habitat types. *I. persulcatus* is more adapted to theropencedrymion, while *H. concinna* is more adapted to herbosa. Our study site Heiniubei (S1) is near a highway, with fewer trees and vegetation and has a higher volume of tourist activity. The other study site, Dashigou (S2), is located in the remote mountains, with denser vegetative cover. These environmental factors may be responsible for the heterogeneous distribution between tick species. This in turn likely contributes to differences in *Babesia* prevalence and species diversity. The prevalence of *Babesia* in two regions had significant difference (χ^2^=4.72, *P*=0.04). Infection with *Babesia* spp. in ticks occurred mainly in the Heiniubei study site, where more *H. concinna* found suitable habitats, such as *Equisetum ramosissimum* grass on concave slope. The significant environmental variables in the logistic regression were vector species, concave slope, and *Equisetum ramosissimum* dominant grass. The concave slope might be appropriate habitat for rodent hosts. *Equisetum ramosissimum* is a hollow, nodulated perennial herb which often grows in humid ground under the trees on the slope. This could be the favorable niche for ticks during hatching and molting during their final developmental stages. Maintenance of *Babesia* in a region ultimately requires some connectivity between vector ticks and hosts as well as human activity in the field. Thus, landscape features that influence population connectivity can inevitably be related to *Babesia* dynamics.

Based on geospatial analysis, we found a cluster distribution of positive ticks for *B. venatorum* and HLJ-242 like variants (Figure 3). This cluster includes all three waypoints where there are positive samples at two continuous years. Positive ticks collected in the cluster (waypoint 1) were 15.36 times (P=0.045) more likely to be infected with *B. venatorum* than those collected elsewhere from the transect. These findings are of importance for the assessment of regional and environmental risks of exposure for human babesiosis in northeastern China. Previous reports on natural foci of *Babesia* often comprised a vastly larger scale on the order of counties or the equivalent. By means of a three-year longitudinal study that mapped the location of ticks testing positive for *Babesia* DNA, we were able to identify small foci of transmission. *Babesia* foci may persist depending on transovarial and transstadial transmission within ticks, as well as stable infection in hosts. As previously reported, reservoir hosts for *B. microti* are small mammals and shrews whereas *B. divergens* and *B. venatorum* are found on large ruminants[32]. Further investigation of the ecology and epidemiology of these seven *Babesia spp*, including information on infection rates in humans and livestock, is needed to better understand the burden of disease in these populations. However, the results of this study could be limited by the small area of investigation, around 3000~4000 square meters (approximately 70 m diameter), small sample size, and the impact of consecutive collecting on tick densities. In addition, we did not consider the habitat-related factors (temperature, humidity, soil composition or chemistry, protozoal fauna) which may serve as the basis for the foci in this study. The economic conditions, population immunity, and immigration may also impact the transmission of the disease. Thus, these factors need to be considered in future studies.

**Figure 3.**
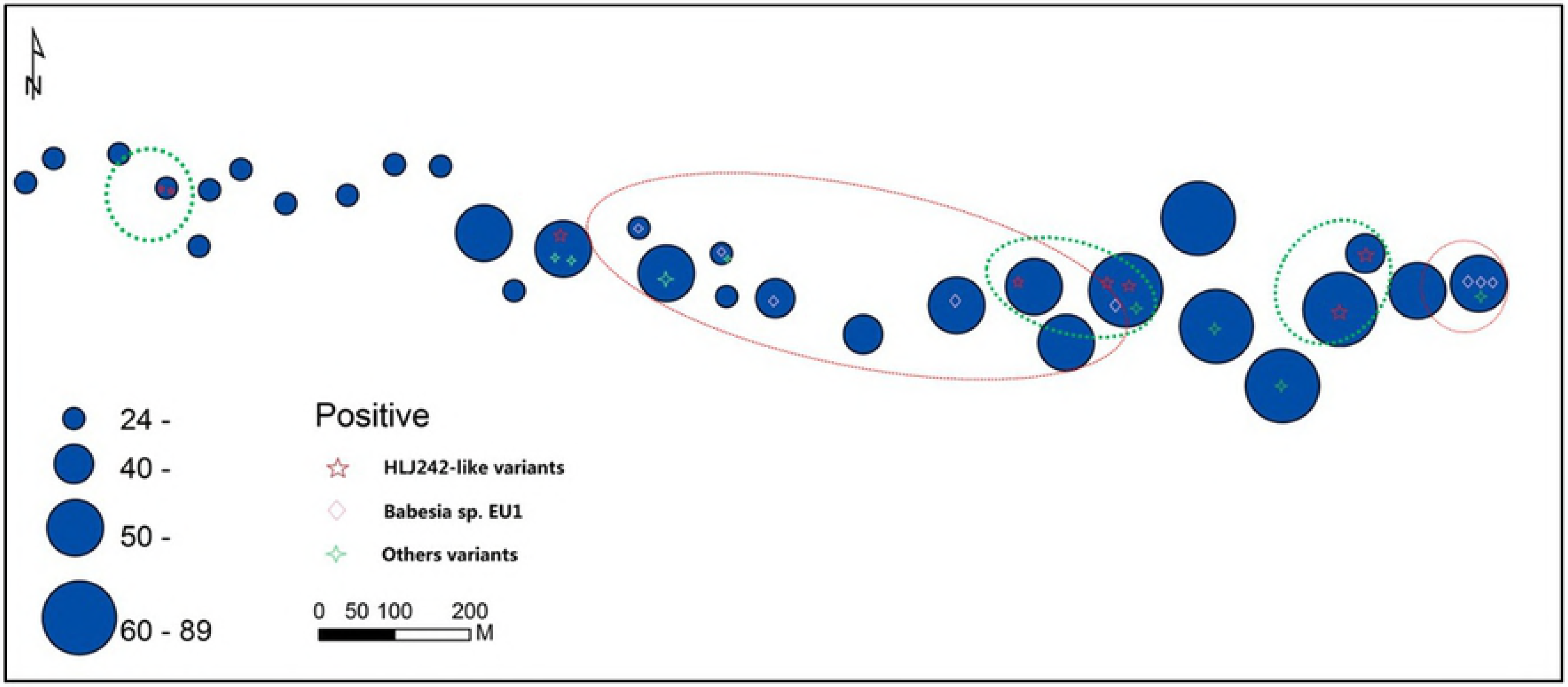
GIS mapping of the each ticks that tested positive predominantly *Babesia* spp. including *Babesia venatorum* and novel variants *Babesia* sp. HLJ242-like strain in this study, and others species by PCR from each waypoint. The size of the dot at each waypoint is proportional to the number of ticks at each site. The location of the cluster for *Babesia venatorum* and HLJ242-like variants is circled.

Our study sites are known to contain tick-borne encephalitis, Lyme disease and spotted fever Rickettsia. This study confirmed that the area also contains *Babesia*, with great diversity within a small natural area. B. venatorum is the predominant species found in *I. persulcatus* for these areas, which is also the main species of *Babesia* recovered from patients in the corresponding areas[14]. The presence of these *Babesia* species, which are of probable medical relevance, in a suburban forest where *I. persulcatus* and *H. concinna* tick density is high requires further attention. Infection by *Babesia* spp. in patients presenting with nonspecific symptoms and history of tick bite or possible tick exposure should be considered, which requires medical awareness of neglected tick-borne pathogens. Whether the two potentially new *Babesia* species identified in this study exist in humans in these areas requires further investigation. Follow-up of the present study is underway to survey the prevalence of *Babesia* spp. in these areas.

## Data accessibility

The18SrRNA gene nucleotide sequences of *Babesia* spp. obtained in this study were deposited in GenBank under accession numbers JQ993417—JQ993430, GU358687, JX542614, JX542615 and KF582564 respectively.

